# Quantifying the economic impact of *Peristenus relictus* establishment on host *Lygus hesperus* populations in California organic strawberry

**DOI:** 10.1101/2022.02.10.480012

**Authors:** Diego J. Nieto, Emily Bick, Charles H. Pickett

## Abstract

*Lygus hesperus* is a key pest in California-grown strawberries, in part because it lacks specialized natural enemies. In response, the specialist European nymphal parasitoid *Peristenus relictus* was released, and subsequently established, along the California Central Coast. A simulation model, based on field-collected *L. hesperus* immigration rates and 16 years of parasitism data, compared simulated pest populations with and without parasitism by *P. relictus*. After 10 years, *L. hesperus* population reductions exceeded 95% in weeds and alfalfa, and 70% in strawberry. These simulated results were in agreement with field-based *P. relictus* longitudinal studies. The averted value loss associated with *L. hesperus*, due to the establishment of *P. relictus*, is estimated at $1,901 per acre in organic strawberry.

## Introduction

*Lygus hesperus* (Hemiptera: Miridae) causes substantial fresh market yield losses in California-grown strawberries by feeding on flowers and small developing fruit, which in turn distorts a strawberry’s appearance. Such feeding damage has been exacerbated by the lack of specialist parasitoids targeting *L. hesperus* nymphs in California strawberry (Van Steenwyk and Stern 1977). In response, a new-association biological control program was initiated and led to the eventual release of the European lygus bug parasitoid, *Peristenus relictus*, on the California Central Coast (Pickett et al. 2009). *Peristenus relictus* populations subsequently dispersed southward, such that they now encompass all major strawberry-producing regions in the state (Nieto et al. 2020).

Upon establishment on the central coast, *P. relictus* demonstrated that it could regulate this pest population, particularly in “stable” environments that were not exposed to insecticide applications (Pickett et al. 2017). Specifically, parasitism by *P. relictus* in these undisturbed environments (e.g., riparian habitat) was significantly correlated with *L. hesperus* nymphal density and eventually reduced pest populations by 99%, relative to pre-release levels. In organic strawberry, *L. hesperus* densities were reduced by 69%, 12 years after the initial release of *P. relictus*. However, creating an economic value that could be associated with such biological control using field studies alone is challenging, especially in crops that experience relatively low pest densities, as is the case with *L. hesperus* in strawberry. Rather, a stage-structured degree day model was utilized to simulate *L. hesperus* populations with and without nymphal parasitism by *P. relictus* in order to determine the economic value associated with these biocontrol services.

## Methods and Materials

### Parasitism data

Parasitism data, which was based on field-collected *L. hesperus* nymphs, informed the biocontrol parameters of *P. relictus* that were used in the simulations. *Lygus hesperus* nymphs were collected in weeds, alfalfa (primarily trap crops) and strawberry fields in Santa Cruz and Monterey Counties from 2002-2018 (excluding 2016). In total, 3,049 samples, using either a sweep net or hand-held vacuum, were collected from 30 sites, resulting in 47,707 collected nymphs and 21,104 dissections.

### Model Structure

To predict *L. hesperus* population dynamics, a stage-structured degree day (DD) model that simulates immigration, reproduction and mortality was developed. The model simulates and records the stage structure for *L. hesperus* (as number per instar) by simulating daily (24-hour) time steps for a duration of 3,652 days (10 years). Insect stages are implemented accounting for a lag between individuals entering and exiting developmental stages (Manetsch 1976), including eggs (Vansickle 1977), in accordance with the distributed delay principal. Immigration was simulated with the daily addition of 7.716 × 10^−5^ ovipositing *L. hesperus* per strawberry plant (Bick 2019). Mean daily temperatures obtained from the Watsonville West II #209 weather station (R Development Core Team 2016, CIMIS 2019) were used to calculate the DDs using the normal model (Duffy et al. 2017). Simulated partial or full individual(s) transition through developmental stages or exit the simulation based on accumulated DDs, which in turn is based on life tables developed by Leigh (1963) and Strong et al. (1970). Oviposition was normally distributed over the DD for all individual or partial individual adults (both males and females), using an adjusted fecundity rate of 105 eggs per adult, given the 1:1.1 sex ratio of adults and the established female fecundity rate of 210 eggs (Leigh 1963, Strong et al. 1970).

### Simulation Experiment

To predict the long-term regulation of *L. hesperus* populations by *P. relictus*, a proportion of 5^th^ instar nymphs were removed from the simulation during the transition between nymphal and adult stages (Fig. 1). This proportion was based on parasitism rates observed in weeds, alfalfa, and strawberries from 2002-2018 (as described above). For each plant type, a parabola was fit to (1) the first quartile, (2) the mean, and (3) the third quartile of field parasitism rates, termed ‘high’, ‘mean’, and ‘low’ parasitism respectively. Plant specific simulations were run pulling daily parasitism rates from the parabolas. Simulations were run for a 10-year timeframe, starting April 1^st^, 2009, and ending in 2019. Results are reported as the proportion of a *L. hesperus* population that is removed due to parasitism over 10 years, when compared with a similar population without such parasitism. An open-source modeling platform called the Universal Simulator was used to run the model. This tool was developed in the Qt integrated development environment and programed with the C++ programming language (Holst 2013). Code and instructions for model replication are found at https://www.ecolmod.org/.

**Figure 1.**
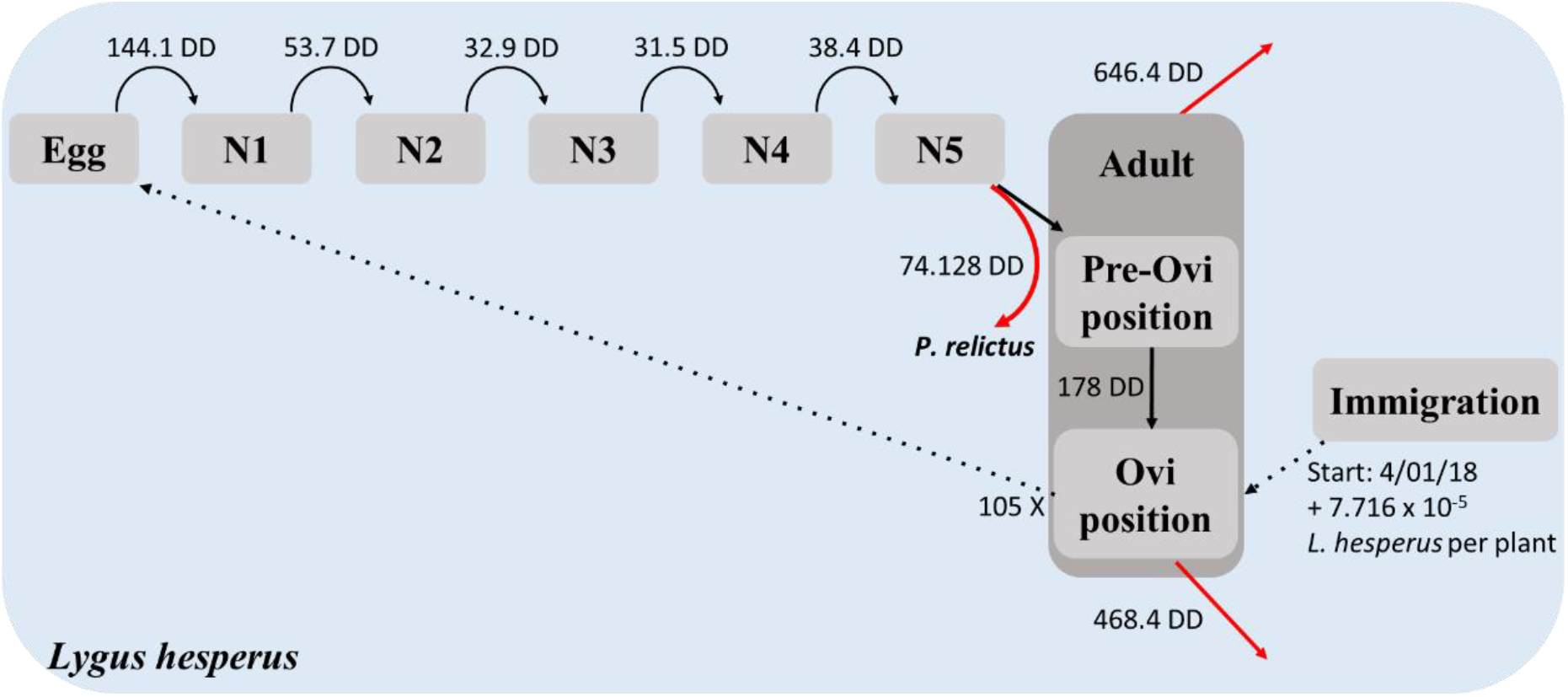
Diagram of stage-structured degree day model predicting *L. hesperus* population dynamics seasonally regulated by *P. relictus*. Black arrows represent movement between stages and red arrows represent exiting simulation.

The economic analysis was conducted by applying the proportion of simulated *L. hesperus* eliminated from their population, as a result of parasitism by *P. relictus*, to a valuation of *L. hesperus* damage. This reduction was then compared with a similar damage valuation that lacked the population reductions that resulted from biological control. This value loss estimate was acquired by correlating organic strawberry damage rates and lygus bug nymph densities over four field seasons in Monterey County (unpublished data). For instance, if damage associated with a fixed pest density was valued at $100, and simulations showed that biocontrol reduced a pest’s population by half, then we estimated the economic impact of biocontrol to be $50. A value loss estimate associated with a *L. hesperus* density of one nymph per 100-suction sample was used in order to produce conservative economic values for *P. relictus*.

## Results and Discussion

Multiyear parasitism averages were generated for *L. hesperus* nymphs collected in weeds, alfalfa, and strawberry (Fig. 2). Parasitism rates of *L. hesperus* were generally greatest in weeds, followed by alfalfa, then strawberry. Relative differences in parasitism between crops is likely attributable to dissimilarities regarding host density and exposure to management efforts. Specifically, low host densities, along with multiple insecticide applications and persistent tractor-mounted vacuuming, contributed to lower parasitism levels in strawberry. Consequently, the simulation for strawberry does not have a “low” parasitism scenario, as the first parasitism quartile was 0%.

**Figure 2.**
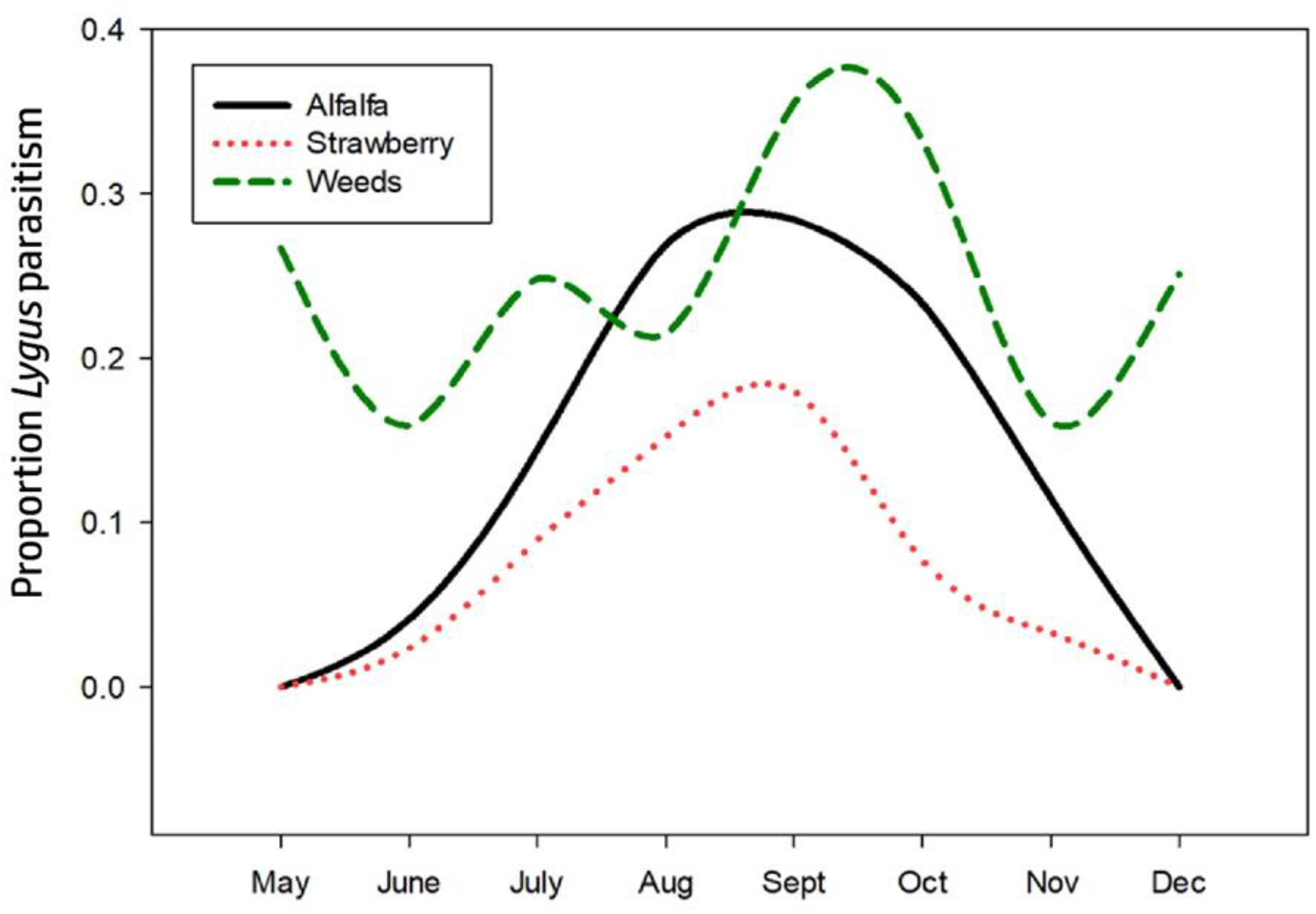
Proportion parasitism of *L. hesperus* nymphs in Santa Cruz and Monterey Counties from 2002-2018.

The simulated proportion of *L. hesperus* nymphs removed from weeds, alfalfa and strawberry due to parasitism by *Peristenus relictus*, when compared with pest population development without such parasitism, is shown in Figure 3. Over 95% of *L. hesperus* populations in weeds and alfalfa were reduced using mean parasitism levels over a 10-year span. Over 70% of the *L. hesperus* population in strawberry was reduced using mean parasitism levels after 10 years. Simulation results from weeds and strawberry approximate the nymphal reductions observed through longitudinal sampling by Pickett et al. (2017): nymphal densities dropped by ∼99% and 69% at two riparian (i.e., weedy) field sites and an organic strawberry ranch, respectively.

**Figure 3.**
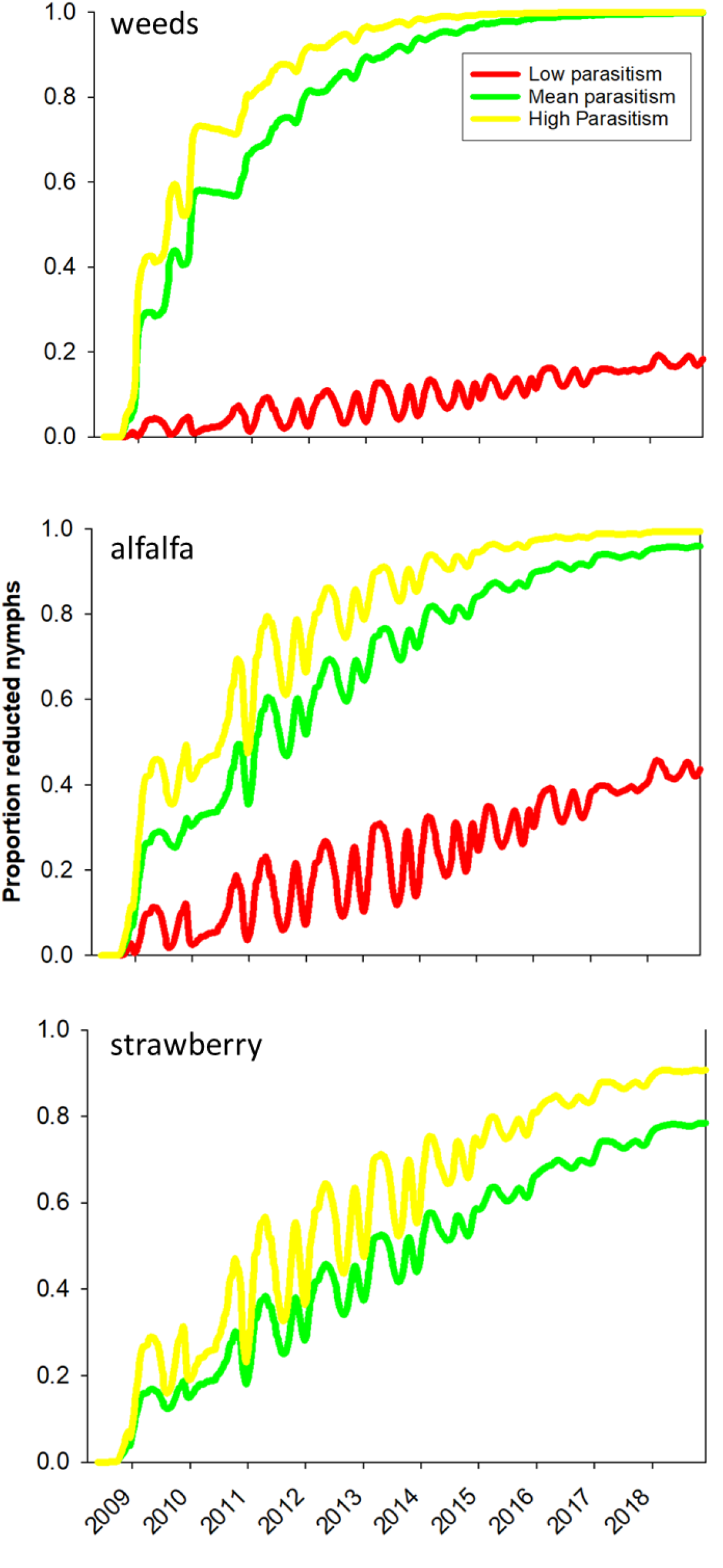
Simulated proportion of lygus bugs nymphs removed due to parasitism relative to predicted population development without parasitism during a 10-year timeframe. Model outputs are shown from weeds, alfalfa, and strawberries under ‘high’, ‘mean’ and ‘low’ parasitism.

Nonetheless, it’s possible that more comprehensive simulations would more accurately confer the biocontrol dynamics in this system. For instance, tri-trophic modeling may better reflect *L. hesperus* productivity (i.e., fecundity), host density, floral resource availability, and potential for parasitoid mortality. Among the many questions that arise: how does spring-time parasitism in weeds affect *L. hesperus* immigration rates? Is this pest more productive in alfalfa than strawberry, and if so, does that negate higher parasitism rates in trap crops? How would a beneficial insectary in strawberry affect pest reductions? Clearly, more research is needed in order to construct and inform increasingly comprehensive biocontrol model simulations.

The baseline value loss estimate for a single *L. hesperus* nymph (resulting from feeding damage) in organic strawberry is $2,715 per acre. When the simulated reduction of this pest’s population (70% after 10 years) is applied, the value loss differential between baseline and biocontrol-added treatments equals $1,901 per acre. When these savings are applied to all 1,973 organic strawberry acres in Monterey and Santa Cruz Counties in 2021, the total averted value loss equals $3.75 million.

This economic analysis provides a loose approximation of the savings provided by *P. relictus*. However, conservative figures were selected at every opportunity in order to reduce the probability of overestimating this parasitoid’s value. Additional field-collected data could bolster confidence levels pertaining to the model’s assumptions. Finally, a return-on-investment ratio should be calculated to demonstrate the worthiness of this biological control program (e.g., foreign explorations, permitting, quarantined colony production, releases, field validations, etc) and help justify similar future efforts.

---

**Table 1.**
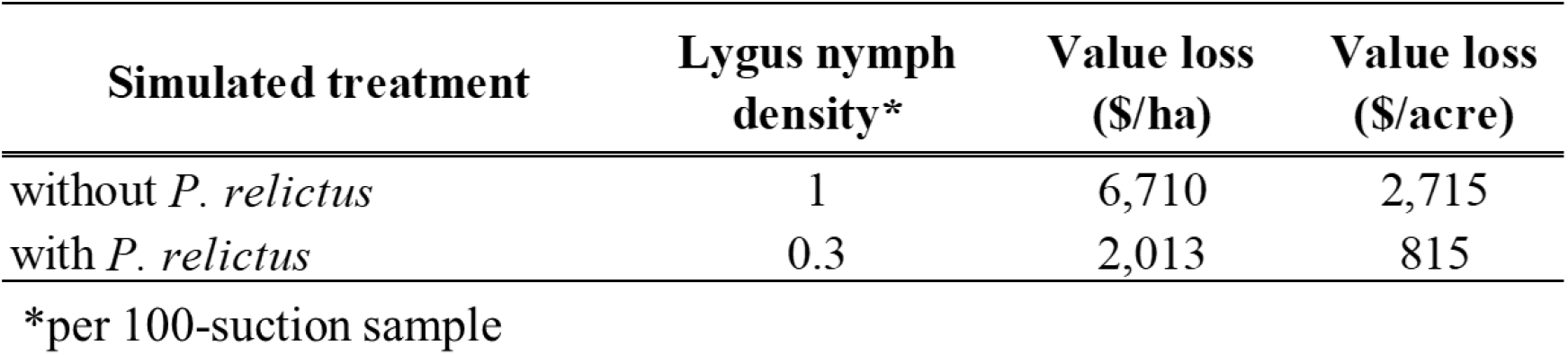
Valuation of lygus bug biological control in California organic strawberry. Treatment-based value differential acquired by applying simulated lygus bug population reductions to corresponding value loss estimates.

| Simulated treatment | Lygus nymph density* | Value loss (\$/ha) | Value loss (\$/acre) |
| --- | --- | --- | --- |
| without *P. relictus* | 1 | 6,710 | 2,715 |
| with *P. relictus* | 0.3 | 2,013 | 815 |

\*per 100-suction sample

## References Cited

Bick, E. N. 2019. Plant-insect Interactions to Promote Pest Management in Strawberry and Water Hyacinth Ecosystems. University of California, Davis.

CIMIS (California Irrigation Management Information System). 2019. CIMIS station reports. https://cimis.water.ca.gov/WSNReportCriteria.aspx.

Duffy, C., R. Fealy and R. M. Fealy. 2017. An improved simulation model to describe the temperature-dependent population dynamics of the grain aphid, Sitobion avenae. Ecological Modeling 354: 140–171.

Holst, N. 2013. A universal simulator for ecological models. Ecological Informatics 13: 70–76.

Leigh, T. F. 1963. Life history of Lygus hesperus (Hemiptera: Miridae) in the laboratory. Annals of the Entomological Society of America 56: 865–867.

Manetsch, T. J. 1976. Time-varying distributed delays and their use in aggregative models of large systems. IEEE Transactions on Systems, Man, and Cybernetics 8: 547–553.

Nieto, D. J., J. Buhler, and M. P. Seagraves. 2020. Documenting the expanded southern range of the introduced parasitoid Peristenus relictus (Hymenoptera: Braconidae) in California. Biocontrol Science and Technology 30: 499–504.

Pickett, C. H., D. J. Nieto, J. A Bryer, S. L. Swezey, and M. Stadtherr. 2017. Post-release dispersal of the introduced lygus bug parasitoid Peristenus relictus in California. Biological Control 114: 30–38.

Pickett, C. H., S. L. Swezey, D. N. Nieto, J. A. Bryer, M. Erlandson, H. Goulet and M. D. Schwartz. 2009. Colonization and establishment of Peristenus relictus (Hymenoptera: Braconidae) for control of Lygus spp. (Hemiptera: Miridae) in strawberries on the California Central Coast. Biological Control 49: 27–37.

R Development Core Team. 2016. R: a language and environment for statistical computing, version 1.0.143. R Foundation for Statistical Computing, Vienna, Austria.

Strong, F. E. and J. A. Sheldahl. 1970. The influence of temperature on longevity and fecundity in the bug Lygus hesperus (Hemiptera: Miridae). Annals of the Entomological Society of America 63: 1509–1515.

Van Steenwyk, R. A. and V. M. Stern. 1977. Propagation, release, and evaluation of Peristenus stygicus, a newly imported parasite of Lygus bugs. Journal of Economic Entomology 70: 66–69.

Vansickle, J. 1977. Attrition in distributed delay models. IEEE Transactions on Systems, Man, and Cybernetics 7: 635–638.

